# Rotavirus RNA chaperone mediates global transcriptome-wide increase in RNA backbone flexibility

**DOI:** 10.1101/2022.01.04.474988

**Authors:** Aaztli Coria, Anastacia Wienecke, Alexander Borodavka, Alain Laederach

## Abstract

Due to genome segmentation, rotaviruses must co-package a set of eleven distinct genomic RNAs. The packaging is mediated by the RNA chaperone NSP2. While the activities of RNA chaperones are well studied with short RNAs, little is known about their global effect on the entire viral transcriptome. Here we used Selective 2′-hydroxyl Acylation Analyzed by Primer Extension and Mutational Profiling (SHAPE-MaP) to systematically examine the secondary structure of the rotavirus transcriptome alone and in the presence of NSP2. Surprisingly, SHAPE-MaP data reveal that despite the well-characterized helix-unwinding activity of NSP2 *in vitro*, its incubation with rotavirus transcripts does not induce a significant change in the SHAPE reactivities. However, a quantitative analysis of the per nucleotide mutation rate, from which SHAPE reactivities are derived, reveals a global five-fold rate increase in the presence of molar excess of NSP2. Further analysis of the mutation rate in the context of structural classification reveals a larger effect on stems rather than loop elements. Together, these data provide the first experimentally derived secondary structure model of the rotavirus transcriptome and reveal that NSP2 exerts a larger effect on stems, while acting by globally increasing RNA backbone flexibility in a protein concentration-dependent manner.

## INTRODUCTION

Virions of rotaviruses (RV), a large group of important human and animal pathogens, contain eleven distinct dsRNA molecules, or genomic segments, all of which are essential for rotavirus infectivity (1–3). Each dsRNA segment serves as a template for transcribing differently sized (0.67-3.4 kb) protein-coding +ssRNA transcripts (mRNAs) that can fold into regulatory structures, such as those predicted to be found in the 5′ and 3′ untranslated regions (UTRs) (4, 5). In addition, consensus sequences found in 5′ and 3′ terminal regions of these segments also have important regulatory roles in viral infection, such as recruiting viral and potentially host proteins to enhance viral translation (6, 7). Li *et al* provided one of the most comprehensive studies of rotavirus RNAs based on the minimum free energy consensus structures deduced from multiple sequence alignments and covariation analysis of group A rotavirus (RVA) strains (5). This study revealed multiple conserved RNA regions present in all RNA segments that were proposed to fold into regulatory elements. However, in this study, structure probing validation of generated models was only carried out for the smallest segment 11 transcript. Structural analysis of a complete rotavirus transcriptome with single-nucleotide precision would enable identification of readily identifiable RNA structures in rotavirus genome that would galvanize their functional validation through recently established reverse genetics studies (8). Structural descriptions of multiple viral RNAs have uncovered myriads of previously unknown functional regulatory motifs (9, 10). Thus, the first step of identifying functional structural domains in viral RNAs is to model experimentally validated RNA secondary structures.

Importantly, rotaviruses encode an RNA chaperone NSP2 that is believed to be involved in RNA selection and stoichiometric segmented genome packaging (3, 11). Single-molecule fluorescence studies of NSP2-mediated remodeling of short (< 50 nts) RNA stem-loops indicate that NSP2 binds single-stranded RNAs with nanomolar affinity, causing transient helix destabilization and melting of its secondary structure (12, 13). These studies also revealed that NSP2 octamers can bind to RNA stem-loops in both open and closed conformations, raising further questions about the mode of action of NSP2 as an RNA chaperone that promotes inter-segment RNA interactions (13). While these studies were important for our understanding of the RNA chaperone-mediated RNA annealing, there is no direct evidence of structural reorganization of the RV RNA upon NSP2 binding. Moreover, NSP2 binding itself poses another major challenge for modelling RV RNA structures, as there is very little understanding of how multivalent RNA chaperones can alter RNA folding on a transcriptome level (14). NSP2 is an RNA chaperone known to melt RNA helices, its helix destabilizing properties have been characterized using single-molecule FRET and gel electrophoresis with synthetic RNA and DNA substrates (12, 13, 15). However, this helix-destabilizing activity has yet to be investigated using RV RNA at a transcriptome scale.

RNA folding is mediated by base-pairing that forms secondary structures which play a myriad of fundamental regulatory roles including gene expression (9, 16, 17). To experimentally interrogate RNA structure, chemical structure probing coupled with thermodynamic modeling provides single nucleotide resolution information with transcriptome scale throughput (18, 19). This is particularly pertinent for viruses, whose RNA genomes, or ssRNA precursors of dsRNA genomes, can be directly translated into viral proteins, or participate in the assembly of a nascent virus (20, 21). Thus, experimentally investigating RNA structure with chemical probing allows us to identify potentially new and important roles for RNA structure in rotaviruses.

Here, we provide experimentally validated structure models of a complete rotavirus RNA pre-genome probed by selective 2′-hydroxyl acylation analyzed by primer extension (SHAPE) and mutational profiling (MaP) analysis (19, 22). We identified multiple conserved structured regions with low SHAPE reactivity located in the 5′ terminal regions of RV RNAs. We show that the RNA chaperone NSP2 binding to RV transcripts dramatically increases mutational rates induced by the SHAPE reagent, but has no effect on the computed SHAPE reactivity, suggesting a global effect on backbone flexibility with minimal structural rearrangement. Analysis of SHAPE-informed secondary structure models in the presence of NSP2 reveals that despite its promiscuous RNA binding mode, its binding preferentially affects base-paired elements, consistent with the RNA chaperone activity. These findings support a model in which promiscuous binding of NSP2 to the rotavirus transcriptome allows this chaperone to uniformly increase the flexibility of the RNA backbone, thus leading to the increased accessibility of RNA towards the establishment of inter-molecular contacts.

## MATERIALS AND METHODS

### Rotavirus Transcript Preparation

In this study we used previously characterized pUC19 clones encoding for segments 1-11 of bovine rotavirus A (strain RF, G6P6[1]) (13, 23). Sequences for the 11 segments are listed in Supplementary File S3 in SNRNASM format. The plasmids coding for segments 2 and 8 were digested using BbsI restriction enzyme (New England Biolab, NEB), the plasmid coding for segment 10 was linearized with BsaI (NEB), and the plasmids coding for the remaining segments were linearized using BsmBI (NEB). Following linearization, the digestion product was purified, and each RV segment was then transcribed using 1 μg of linearized DNA with HiScribe T7 from NEB at 37°C for 3 hours. The transcription reaction was subsequently treated with Turbo DNAse (ThermoFisher) for 30 minutes at 37. The RNA was purified using Purelink RNA columns (ThermoFisher) and eluted in 30 μL of nuclease free water. Each RNA segment quality and size were assessed on a 2% denaturing MOPS formaldehyde agarose gel, prior to loading on the gel RNA samples was denatured at 70°C in 50% (v/v) formamide and resolved on a denaturing formaldehyde-MOPS 1% agarose gel, as described previously (24).

### NSP2 Expression and Purification

A sequence-verified pET-28b-NSP2 construct was used for protein expression in BL21(DE3) *E*.*coli* as described in (15, 25). Ni-affinity-purified NSP2 fractions were further purified over a HiTrap SP cation-exchange column (Cytiva), and the concentrated peak fractions were resolved on a Superdex 200 10×300 GL column and pre-equilibrated with RNAse-free SEC buffer (25 mM HEPES-Na, pH 7.5, 150 mM NaCl) to ensure high purity and homogeneity of the preparation. The peak fraction was at least 99% pure by SDS-PAGE, with a characteristic 260nm/280nm absorbance ratio of < 0.57 suggesting that the purified protein was pure from contaminating nucleic acids.

### Structure Probing using SHAPE-MaP

To prepare RV RNA for structure probing – for single RNA experiments, each segment-specific transcript was diluted to a final concentration of 260 nM and incubated at 65°C for 5 minutes, then slowly cooled back to room temperature for 30 minutes and kept at 4° C. For transcriptome-wide experiments, each RV transcript was diluted to a concentration of 25 nM, incubated at 65°C for 5 minutes, then slowly cooled back to room temperature. Eleven distinct transcripts were then pooled, resulting in an equimolar mix of all 11 genomic RV transcripts. NSP2 protein sample was diluted in 10 mM HEPES-Na pH 7 to concentrations required for the titrations described: i.e., 40 μM, 20 μM and 10 μM. The RNA sample was then buffered in the following conditions: 1 mM MgCl_2_, 200 mM NaCl, and 200 mM HEPES (pH 7). NSP2 was added at an equal volume to the RNA, while for the no protein condition only 10 mM HEPES-Na buffer was added at an equal volume to the RNA sample. The RNA:NSP2 samples were then incubated at 37°C for 30 minutes prior to probing.

For probing, RNA:protein mixes were incubated with 10% (v/v) of 125 mM 5-nitroisatoic-anhydride (5NIA, Sigma) dissolved in dimethyl sulfoxide (DMSO) to achieve a final concentration of 12.5 mM 5NIA. Untreated samples were incubated with 10% DMSO. Both modified and untreated samples were rapidly mixed and then incubated at 37°C for 5 minutes. RNA was subsequently purified using the Qiagen MinElute columns and eluted in 30 μL of nuclease free water. DMS probing was achieved as described in (26), with following modifications: samples were treated with dimethyl sulfate (DMS, Sigma) at a final concentration of 1% DMS in 10% ethanol (v/v), or 10% ethanol for control samples, incubated for 5 minutes at 37°C. RNA was subsequently purified using the Qiagen MinElute columns and eluted in 30 μL of nuclease-free water.

### Library Preparation and Next Generation Sequencing

First stand cDNA synthesis of treated RNA was carried out as previously described in (22). Briefly, 29 μl of purified modified RNA was added to 100 ng Random Primer 9 (NEB) and 0.2 mM of each dNTP. This mix then incubated at 65°C for 5 min then incubated on ice. This reaction was buffered for a final concentration of 50 mM Tris (pH 8.0), 75 mM KCl, 10 mM DTT, and 6 mM MnCl_2_, and 2 μl of SuperScript II Reverse Transcriptase (Invitrogen) was added. The reaction was supplemented with 20 U of Murine RNase inhibitor (NEB) and incubated at 25°C for 10 min, then 42°C for 3 h, followed by 70°C for 15 min. First strand cDNA sample was desalted using G50 columns (GE), and immediately introduced into a second strand synthesis reaction using mRNA Non-Directional Second Strand Synthesis module (NEB). Double stranded DNA was then cleaned up using Ampure XP purification beads and eluted in 30 uL of nuclease-free water, quantified using a Qubit fluorimeter (Invitrogen). Double stranded DNA was fragmented, repaired, and adaptor ligated for sequencing library preparation using the NEBNext^®^ Ultra™ II DNA Library Prep Kit for Illumina^®^. Library quality was assessed using Qubit dsDNA HS Assay (Invitrogen) following the High Sensitivity Bioanalyzer protocol (Agilent). Libraries were then loaded on an Illumina MiSeq at 10 pM and run at 300 cycles (150×2) or 600 cycles (300×2).

### SHAPE-MaP Analysis

Sequencing reads produced from our mutational profiling experiments were analyzed using the ShapeMapper2 pipeline (27), version 2.1.4 using Bowtie2 version 2.3.4.3 as the aligner. ShapeMapper2 computes mutation rates for both treated and untreated samples by counting the mutations at each individual nucleotide along the transcript in comparison to the reference sequence (Supplementary File S3). Reactivity is computed by subtracting the mutation rate of the untreated sample from the mutation rate of the treated sample. Statistical outliers are then excluded from further analysis and reactivities are normalized as previously described (28).

### RNA Secondary Structure Modeling and Forgi Classification

All minimum free energy diagrams were generated using RNAStructure (29) by incorporating SHAPE reactivities as a pseudo free energy term and using a maximum pairing distance of 150 nucleotides unless otherwise stated. Using connectivity tables (.CT) generated from RNAStructure we then used the Forgi python library to classify each nucleotide into a specific RNA secondary structure motif (30). These nucleotide classifications were generated from our SHAPE-informed secondary structures derived from structure probing experiments of the RV transcriptome incubated together at an equimolar ratio.

### Statistical Analysis

Mutation rate distribution analysis and establishment of linear relationships between NSP2 titration points (Figures 3-5) was analyzed using R studio (version 2021.09.1+372)and visualized with the ggplot2 package (31). Box plots and histograms (Figures 5 and 6) comparing the log2FoldChange of RNA structural motifs was done using the python package MatPlotLib.

### Data Availability

Raw sequencing reads are available on the NCBI SRA database under BioProject ID PRJNA791398. All SHAPE data used in this study are available in Supplementary File S3 in SNRNASM format.

## RESULTS

### SHAPE-MaP derived structural analysis of rotavirus A transcriptome

Recently established reverse genetics approaches for rescuing viable rotaviruses utilize RVA transcripts produced by T7 polymerase-driven transcription (8), suggesting that a complete set of eleven non-capped RV transcripts generated by T7 polymerase is sufficient to initiate a replicative cycle. Therefore, we utilized a set of plasmids used for the virus rescue to produce eleven RVA transcripts in vitro (23). The produced RNA was intact and pure to be used for subsequent analyses by high-throughput selective 2′-hydroxyl acylation analyzed by primer extension (SHAPE) to interrogate its secondary structure (Supplementary Figure S1A). To develop an accurate model of RNA structure of the RVA transcriptome, we co-incubated an equimolar mix of eleven RV transcripts (0.25 μM total RNA concentration) in the RNA folding buffer (100 mM NaCl, 100 mM HEPES-Na, 0.5 mM MgCl_2_, pH 7) at 37°C for 30 min. Under these conditions, eleven RVA transcripts did not form stable inter-molecular contacts in our previous single-molecule fluorescence studies (13). The RNA mix was then incubated with the SHAPE reagent 5-nitroisatoic anhydride (5NIA), as described in Materials and Methods (32). Conformationally flexible nucleotides exhibit high reactivities towards the electrophile 5NIA (32), thus allowing us to assess the RV transcriptome reactivity with single-nucleotide resolution. We performed at least two biological replicates of all SHAPE experiments, and SHAPE reactivities for these replicates were reproducible and highly correlated with R^2^ > 0.7 in all cases (Supplementary Table S1). We quantified relative flexibility of each transcript using median SHAPE analysis, as previously described (33–36) (Figure 1A). Regions of low median SHAPE are generally more likely to fold into a single, well-defined conformation, while nucleotides with above median SHAPE generally adopt multiple structures (34). We modeled the secondary structure and calculated base-pairing probabilities for the entire RV transcriptome. To do so, we used our experimentally determined SHAPE reactivities averaged over both replicates as a pseudo-free energy term, as previously described (34, 37, 38). The resulting modelled base-paired nucleotides are shown as arcs, where the color of each arc represents the likelihood of the base-pair formed (Figure 1A). Representative models of regions of low SHAPE (minimum free energy structures) are shown in Figure 1B.

**Figure 1.**
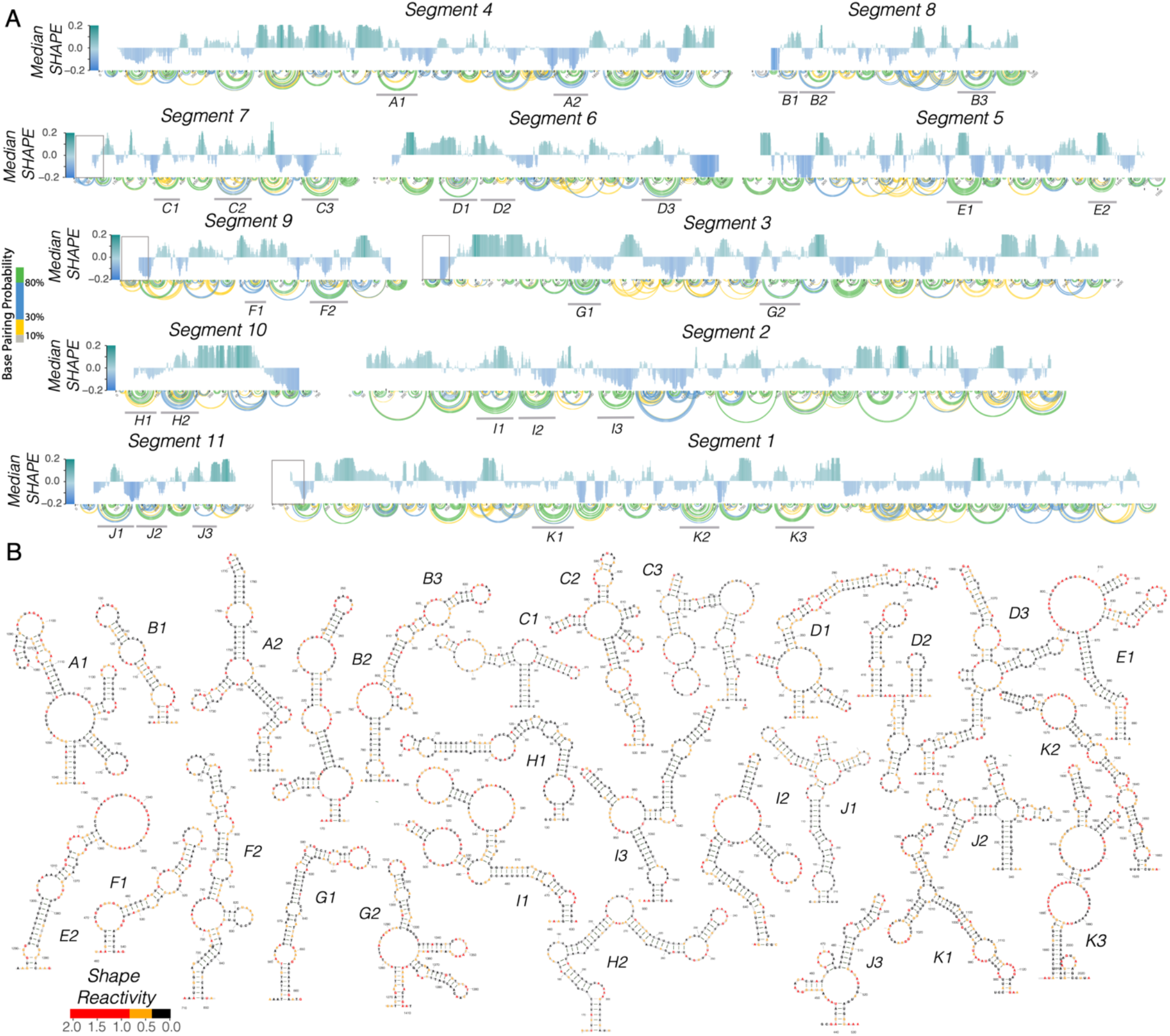
The Rotavirus transcriptome exhibits a diverse assortment of RNA structures. (A) Median SHAPE reactivity of each of the 11 RNA transcripts representing the rotavirus transcriptome averaged over two highly correlated (R^2^ > 0.9) replicates. SHAPE experiments were carried out by incubating equimolar amounts of each RV segment and probed with 5NIA as described in Materials and Methods. A 50-nucleotide window was used to calculate median SHAPE values along each transcript and normalized to the overall median of the individual transcript. Areas of high median SHAPE (i.e., above zero, shown in green) represent conformationally flexible RNA regions. Areas of low median SHAPE (i.e., below zero, shown in blue) represent regions with high propensity to base-pair. Arc plots show pseudo-free energy derived base-pairing probabilities computed using nearest neighbor RNAStructure parameters constrained by incorporating SHAPE reactivities (18, 29). Probabilities of base-pairing are color-coded, with green arcs corresponding to the most probable base pairs, and grey being the least probable ones. Boxed regions indicate conserved nucleotides previously found through conservation analysis (5). (B) Minimum Free Energy (MFE) diagrams of SHAPE informed representative structures within the RV transcriptome (A-K as denoted in panel A). MFE models were generated using RNAStructure parameters as described above.

To further validate our SHAPE-MaP probing data, we compared our SHAPE-informed secondary structure model of the segment 11 transcript to a predicted secondary structure model derived from previously published RNA-RNA SELEX data (13). The analysis revealed that both models were broadly similar (PPV of 66.8% and sensitivity of 84.52%, Supplementary Figure S1B). In addition, we compared our SHAPE-derived models with the secondary structure models derived from the sequence conservation analyses (5). Remarkably, both conserved 3′ terminal regions of segments 10 and 6 exhibited low SHAPE reactivities, consistent with the predicted highly base-paired structures (Figures 2A and B, respectively). We further identified two previously predicted stem loops located in the 3′ terminal region of segment 6. This region has translation enhancing activity important for VP6 expression (39).

**Figure 2.**
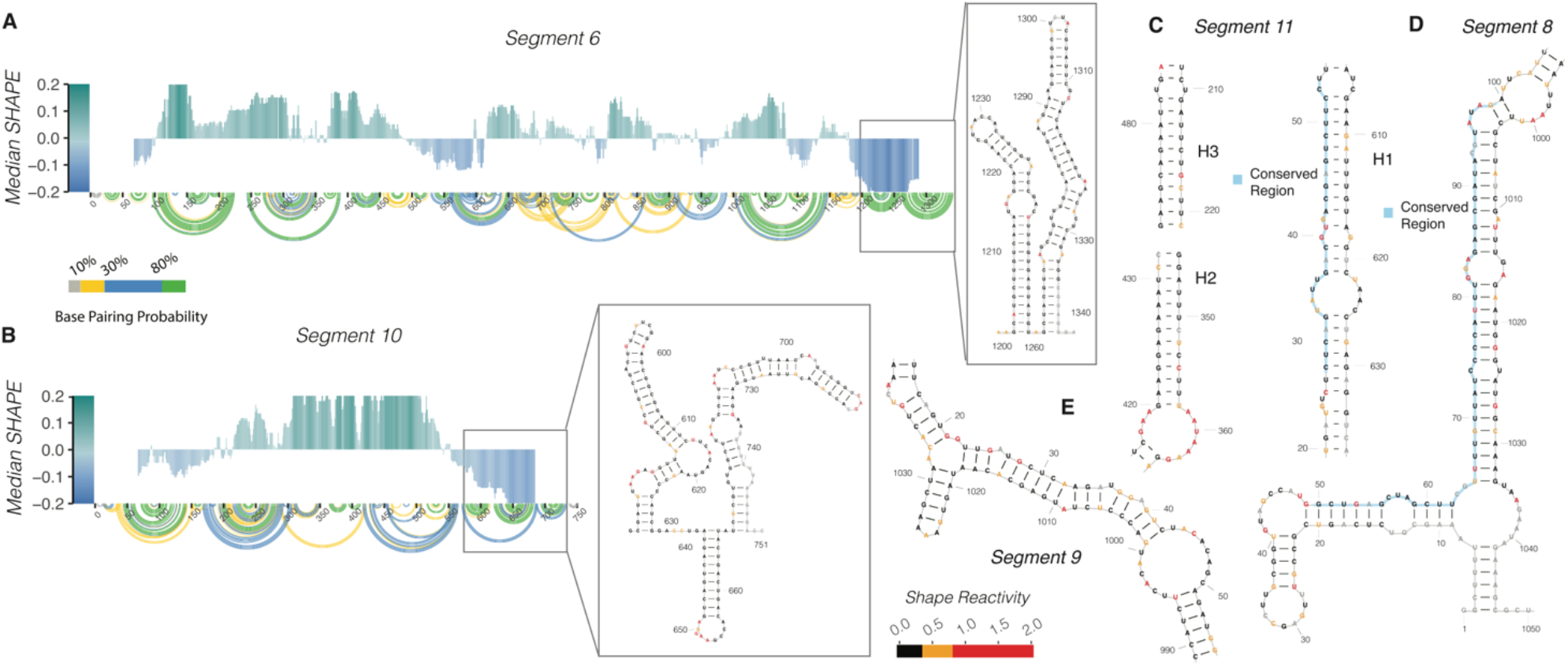
SHAPE-MaP derived structures support previously predicted structures. (A) Median SHAPE reactivity of segment 6 when incubated with all 11 RV segments. Median SHAPE profile and base pairing probabilities were calculated as described in Figure 1. Grey box indicates two terminal stem loops which were previously predicted and were present in our SHAPE informed secondary structure model (5). (B) Median SHAPE-MaP profile and base-pairing probabilities of segment 10 when incubated with all 11 RV segments. Median SHAPE profile and base pairing probabilities were calculated as described in Figure 1. Grey box highlights three stem loops in 3′ terminal region of segment 10 as previously predicted (5). (C) Helix3 (H3) and Helix 2 (H2) were previously described by Li *et al*., and are supported by our SHAPE derived secondary structure model. Highly conserved nucleotides, nucleotides with low normalized mean pairwise distance (nMPD), were previously predicted to participate in LRIs between terminal regions and our SHAPE informed MFE model of segment 11 validates these previous predictions. (5, 40) (D) Segment 8 MFE model calculated by using SHAPE reactivities as a pseudo free energy term. We found segment 8 highly conserved nucleotides (50-98), are located within two connecting stems with two accurately predicted interior bulges as previously proposed by Li *et al*. (5).

Similarly, our SHAPE-MaP data confirmed the presence of long-range interactions (LRIs) previously detected in segments 8, 9 and 11 (Figures 2C, D and E) through conservation and covariation analysis (5). Interestingly, our SHAPE-MaP data suggested a predominant LRI structure amongst two alternative models that had been previously identified in segment 9 by Li *et al* (Figure 2E).

Collectively, these data reveal all RV segments have structured 5′ terminal regions, expanding into coding regions, except for segments 2 and 5. Previously, it was found that the first 78 nucleotides in segment 1 and segment 7 are highly conserved (5). Additionally, the first 82 nucleotides and the first 94 nucleotides of segment 9 and segment 3, respectively, are conserved. These regions of high conservation correlate with regions of low median SHAPE, structured regions, identified in these data (Figure 1A, as indicated by boxes). In contrast, segments 2 and 5 both have unstructured 5′ terminal regions, additionally, neither of these segments exhibit sequence conservation in the 5′ terminal regions, as previously suggested by Li *et al* (5). These data reveal that RV RNA structural elements are found within areas of high sequence conservation, and demonstrate that the previously determined co-variation models are in good agreement with our experimental data.

### Mutation rate analysis reveals that NSP2 increases segment 11 RNA backbone flexibility

To measure the effects of NSP2 on RV transcripts, we began by incubating *in vitro* transcribed segment 11 RNA (0.67 kb) with increasing amounts of the RNA chaperone NSP2, and carried out SHAPE chemical probing (Figure 3A). Individual SHAPE reactivities of nucleotides across the segment 11 transcript are represented in a bar chart, where high SHAPE reactivities above 0.7 (shown as red bars) indicate flexible nucleotides, while nucleotides with low SHAPE reactivities below 0.5 (shown as black bars) represent conformationally constrained nucleotides. Visual inspection of these data does not reveal any obvious large differences in the SHAPE profiles. This is confirmed in Figure 3B, where we plot the SHAPE reactivities of segment 11 alone (x axis) and segment 11 SHAPE reactivities incubated with increasing amounts of NSP2 (y axis). The analysis shows that the slopes for all the titration points were close to 1, and the data remained highly correlated with R^2^ = 0.78 for the 20 μM NSP2 titration end point (Figure 3B, purple line), which is within expected replicate-to-replicate variability (Supplementary Table S2). Further analysis of SHAPE reactivities of segment 11 RNA in the presence of increasing concentrations of NSP2 suggests that the global median of SHAPE reactivities remains unchanged (Figure 3C). Thus, SHAPE-MaP analysis suggests that segment 11 SHAPE reactivities are highly similar in the absence or presence of NSP2, with only very minimal structural rearrangement and no apparent global change in SHAPE reactivity.

**Figure 3.**
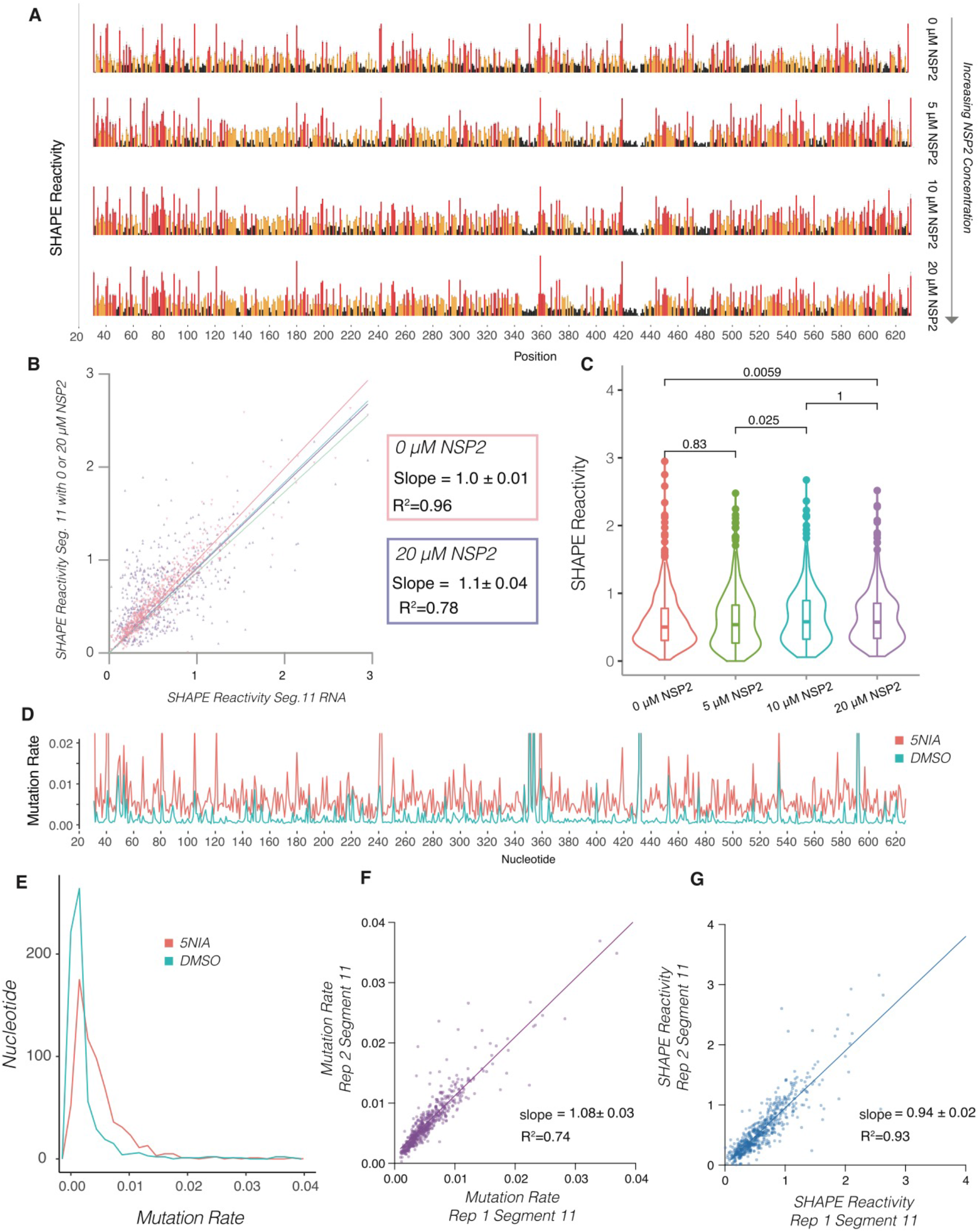
NSP2 addition does not result in detectable changes in the median SHAPE reactivities of segment 11 RNA. (A) Segment 11 was incubated with increasing amounts of NSP2 (0 μM, 5 μM 10 μM, and 20 μM) then probed with 5NIA, as described in the Materials and Methods section. SHAPE reactivities for each of the listed titration points shows no difference in SHAPE reactivity due to the normalization protocol. (B) Scatterplots showing the linear relationship between segment 11 SHAPE reactivity (x axis) and segment 11 SHAPE reactivity when incubated with increasing amounts of NSP2 (y axis). Note high correlation between the data (R^2^ = 0.96, R^2^ = 0.78) and the linear relationship exhibits no significant change between the titration points. (C) Violin plots comparing distribution of SHAPE reactivities of segment 11 RNA upon incubation with NSP2. Boxes represent the 25th/75th interquartile range, and medians are shown as central bands. Addition of 20 mM NSP2 changes the distribution of SHAPE reactivities, while the median SHAPE reactivity does not significantly change, as assessed by the Kruskal-Wallis test (p < 0.05). Note there is no significant difference between the conditions despite a significant global increase in mutation rate. (D) Raw mutation rates of segment 11 transcript probing experiment. The red line is representative of segment 11 when incubated with 12.5 mM of the modifying reagent, 5-nitroisatoic anhydride (5NIA), and the blue is the measured mutation rate across the segment 11 transcript when incubated with the negative control, dimethyl-sulfoxide (DMSO). Normalization of these mutation rates result in the SHAPE profiles shown in panel A. (E) Histogram representation of distribution of nucleotide mutation rates in treated and untreated samples of segment 11 shown in panel D. (F) Scatterplot showing linear relationship of mutation rate data between reproducible replicate experiments. The slope when comparing both replicates is near 1, showing that mutation rate data is highly correlated (R^2^ = 0.74, slope = 1.08 ± 0.03). (G) Scatterplot showing linear relationship of SHAPE-MaP data between reproducible replicate experiments. SHAPE-MaP data are highly correlated (R^2^ =0.93, slope = 0.94 ± 0.02).

SHAPE reactivities are calculated from raw mutation rates (Figures 3D and Figure 3E). The mutation rate of the untreated sample (Figure 3D) is subtracted from the mutation rate of the treated sample (Figure 3D, 5NIA) for each individual nucleotide. Statistical outliers are removed, and the remaining mutation rate intensities are normalized so all reactivities fall between 0 (highly constrained) to 2 (highly flexible) (27, 28). Thus, if NSP2 binding uniformly increases reactivities of all bases, the apparent SHAPE reactivities would remain unchanged, while the mutation rates would uniformly increase for all bases. To test our hypothesis, we investigated the raw mutation rates. Indeed, raw mutational rate analysis confirmed that the distribution of mutation rate for the 5NIA-treated sample is shifted to the right compared to the untreated control sample. Furthermore, we discovered if replicates are carefully executed using a fixed amount of 5NIA reagent, mutation rate replicates (Figure 3F) are highly correlated and have a slope near one, like the SHAPE data (Figure 3G). We therefore hypothesized that if NSP2 has a global effect on backbone flexibility, it would effectively be normalized out using standard SHAPE-MaP data analysis, and only an analysis of the raw mutation rates could reveal any effects.

Remarkably, the addition of NSP2 resulted in a concentration-dependent increase in the distribution of nucleotide mutation rates (Figure 4A), in agreement with our hypothesis.

**Figure 4.**
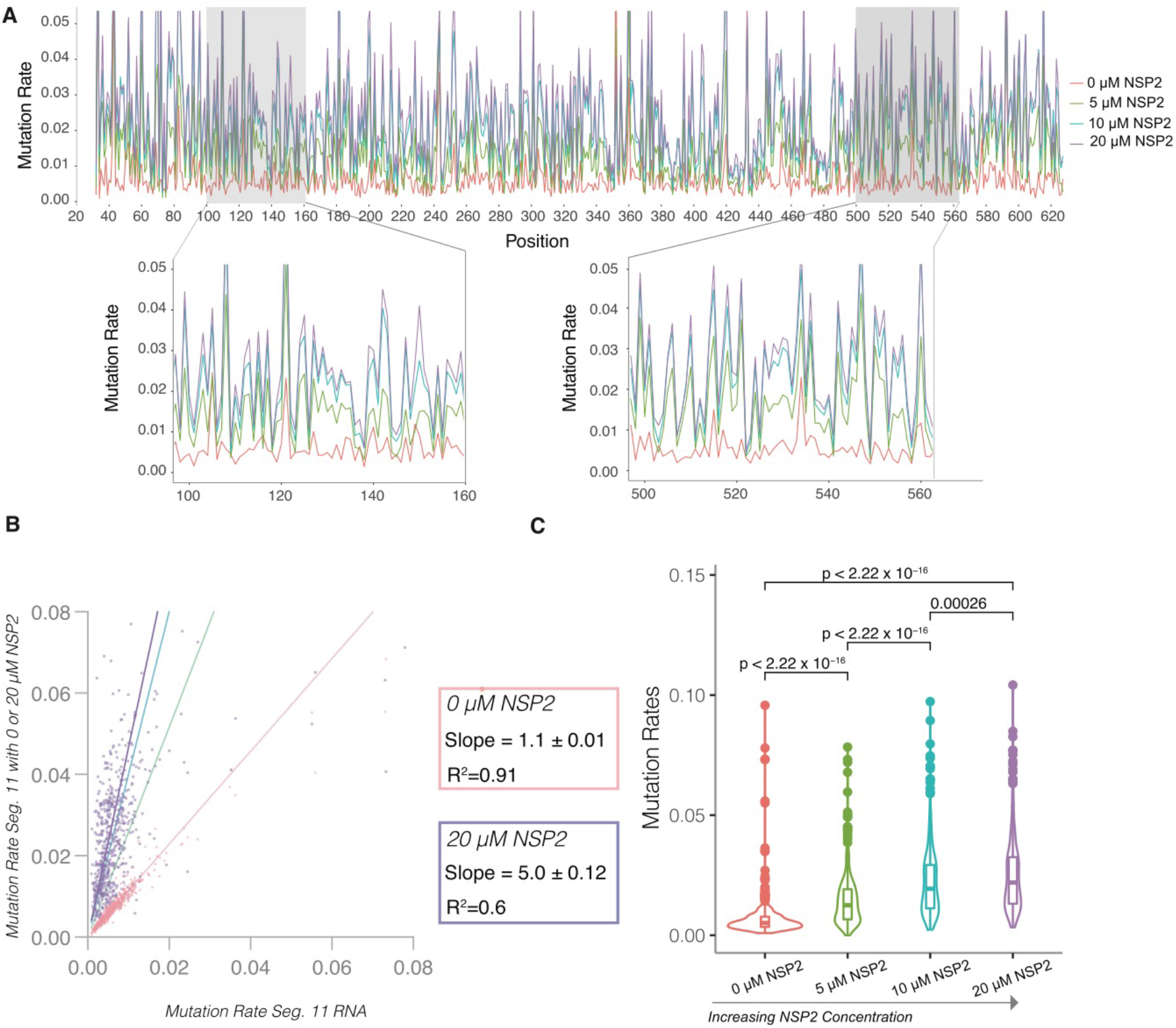
NSP2 uniformly increases segment 11 RNA mutation rate in a concentration dependent manner. (A) Segment 11 was incubated with increasing amounts of NSP2 (0 μM, 5 μM 10 μM, and 20 μM), then probed with 5NIA, as described in the Methods. Higher mutation rates reveal increased flexibility in the RNA backbone, while lower mutation rates indicate less conformationally flexible nucleotides. Note a uniform increase in mutation rates proportional to NSP2 concentration. Zoomed-in: representative regions of segment 11 RNA. (B) Violin plots comparing distribution of mutation rates of segment 11 RNA upon incubation with NSP2. Boxes represent the 25th/75th interquartile range, and medians are shown as central bands. Note, both change in mutation rate distribution and significant change in median mutation rate. Significance values were calculated using Kruskal-Wallis test (p < 0.05) (C) Mutation rates of segment 11 RNA (x-axis) plotted against mutation rates of segment 11 RNA in the presence of increasing amounts of NSP2 to reveal linear relationship with distinct slopes. The slope increase as NSP2 is titrated in, while the linear dependence persists, suggesting a global change in the RNA flexibility in NSP2 concentration-dependent manner.

The pattern of the mutation rates for all bases however remained similar (Figure 4A), which explains a very similar pattern observed in the normalized SHAPE reactivities (Figure 3A).

When we plotted the mutation rate upon addition of NSP2, we observed a strong linear correlation for increasing concentrations of the RNA chaperone, with the slope increase up to 5.0±0.1 for the highest 20 μM NSP2 concentration tested (Figure 4B). We did not attempt higher NSP2 concentrations as those have been previously shown to cause severe RNA aggregation rather than promote strand-annealing reactions (13). This is also consistent with the decreased yields of the recovered RNA from samples incubated with higher amounts of NSP2 that caused protein-RNA aggregation.

The mutation rate increase in the presence of NSP2 was further confirmed by the analysis of the distribution of mutation rates. The median mutation rate increases with increasing NSP2 concentration in a statistically significant way (Figure 4C). Thus, NSP2 has a global effect on segment 11 backbone reactivity as measured by mutation rate, that is masked upon normalization when SHAPE reactivities are computed using the standard normalization procedure. Specifically, we discovered that the quartile normalization step used when deriving SHAPE reactivities from mutational profiling data (28, 33) has the unintended effect of masking these global changes in reactivity in the data.

Finally, to ensure that the changes in mutation rate are mediated by NSP2, and not perturbations caused by the addition of protein, we incubated segment 11 with 20 μM bovine serum albumin (BSA) and found no changes in mutation rate or SHAPE reactivity (Supplementary Figure S2), suggesting the effect we observe here is specific to NSP2.

### NSP2 uniformly increases backbone flexibility in RV segments 5, 6, and 10

To assess if NSP2 increases backbone flexibility across different RV segments, or if the global increase in backbone flexibility is specific to segment 11, we examined *in vitro* transcribed segment 5, 6, and 10 with increasing concentrations of NSP2. We found a similar increase in mutation rate in segments 5, 6, and 10 as segment 11. As can be seen in Figures 5A-C, the mutation rate at 20 μM NSP2 yields slopes > 4 for segments 5, 6, and 10, respectively. Collectively, these data suggest NSP2 increases mutation rate across all segments indiscriminately.

**Figure 5.**
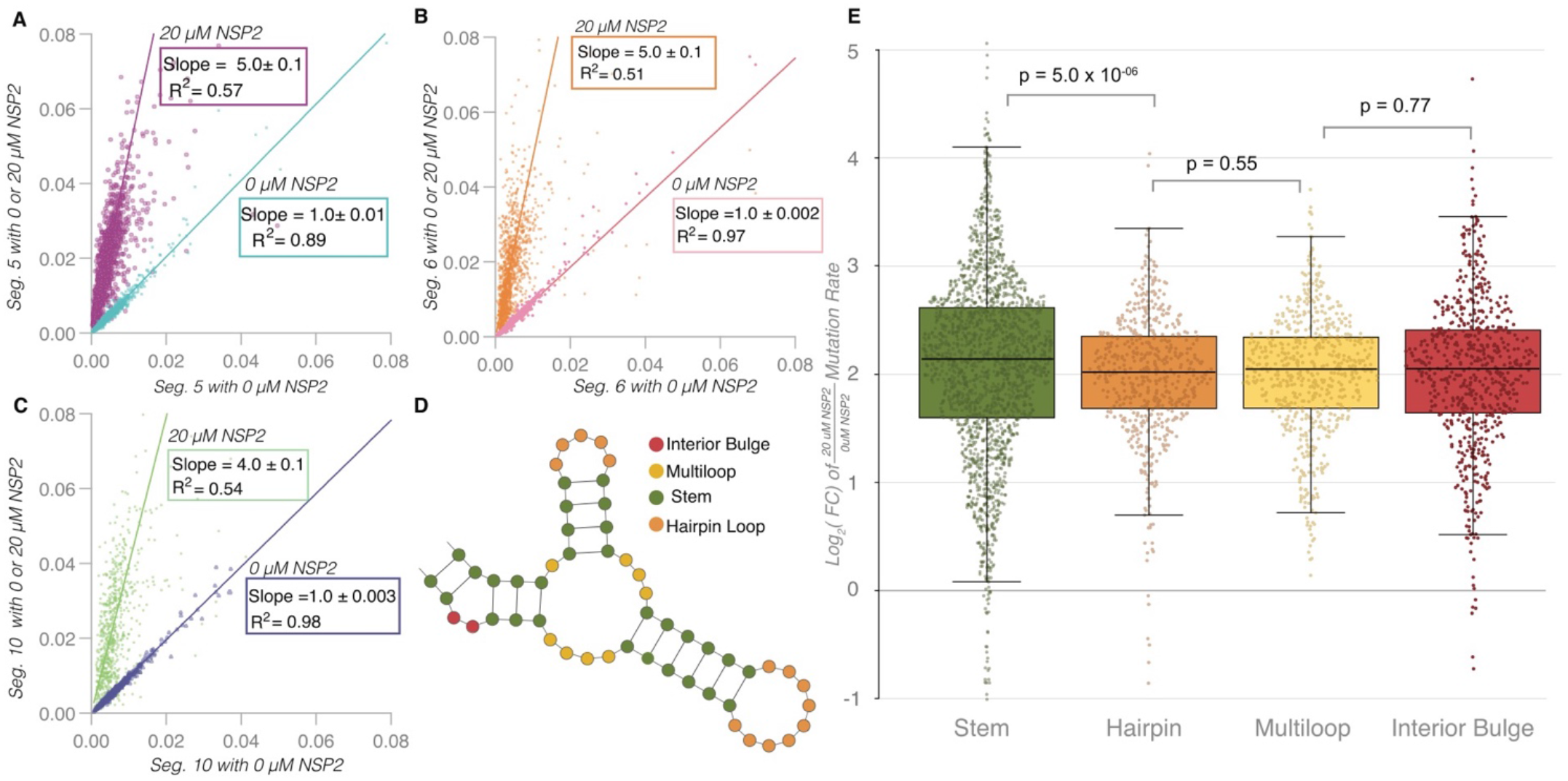
NSP2 uniformly increases backbone flexibility in segment 5, 6 and 10 RNAs. (A-C) Mutation rates of individual RNAs (segment 5, segment 6, and segment 10, respectively) alone (x axis) plotted against the mutation rates of the RNAs incubated with 20 μM NSP2 (y axis). (D) Schematic of structural motifs found within an RNA. Using SHAPE-MaP-informed secondary structure models generated as shown in Fig.1, individual nucleotides (color-coded) were categorized into each of the different structural modalities described: interior bulge (red), multiloop (yellow), stem (green), and hairpin loop (orange) using the Forgi library (Materials and Methods). (E) NSP2-mediated mutation rate change analysis for individual structural motifs as described in panel D. Log_2_(FC) denotes log_2_(mutation rates induced by 20μM NSP2 divided by the mutation rates without NSP2). A vast majority of log_2_(FC) values are positive. Nucleotides involved in stem formation have a higher average log_2_(FC) than those located within predicted hairpins, multiloops and interior bulges. Significance values were calculated using Kruskal-Wallis test.

### NSP2 preferentially increases mutation rate in base-paired structural motifs

The overall pattern of mutation rates remains highly correlated upon addition of even 20 μM NSP2 (Figures 5A-C) for each segment, as does the SHAPE data. As such we can conclude that NSP2 does not have a major effect on the secondary structure of these RNAs even at large molar excess. We were curious to know if we could use mutation rate analysis to determine if NSP2 has any structural specificity at transcriptome scale. We therefore categorized nucleotides of segments 5, 6, 10 and 11 using the Forgi library classifications (interior bulge, multiloop, stem, and hairpin loop) as described in the methods (Figure 5D). These classifications are based on the SHAPE-directed minimum free energy secondary structures reported in Figure 1.

To compare nucleotides in the four different structural contexts, we computed the log_2_(FC) of the mutation rate at 20 and 0 μM NSP2, respectively. As can be seen in Figure 5E, a vast majority of nucleotides have log_2_(FC) > 0, consistent with the positive slopes we observed in the scatter plots in Figures 5A-C. However, we do observe a statistically significant increase in the log_2_(FC) for paired nucleotides (stem) compared with unpaired (hairpin loop, multi-loop, and interior bulge, Figure 5E). The advantage of computing log_2_(FC) in this way is that it allows quantitative comparison of nucleotides with differing mutation rates. Thus, our data show that although the effect of NSP2 on mutation rates is uniformly positive and does not change the overall pattern of SHAPE reactivity (which is indicative of structure), NSP2 has a larger relative effect on the flexibility of stems vs. unpaired nucleotides.

### Structural consequences of whole rotaviral transcriptome on log_*2*_(FC)

As we have previously shown, at sub-nanomolar concentrations, RV transcripts do not appear to interact intermolecularly when NSP2 is absent (13). Indeed, when we compared segments probed alone vs. those probed in the presence of all 11 segments (Supplementary Figure S3), we observed highly correlated mutation rates and slopes near 1. We were therefore interested to see if our log_2_(FC) analysis can detect any statistical differences at transcriptome scale if we incubated all eleven segments with NSP2. We probed the entire transcriptome in the presence of increasing concentrations of NSP2. As can be seen in Figures 6A-D, the 20 μM NSP2 mutation rates are highly correlated in both the absence (x axis, individual transcripts) and presence (y axis, transcriptome) of the other segments, with slopes near 1 in all cases. We also computed slope and R^2^ for increasing concentrations of NSP2 (Table 1) to show that all transcripts exhibit a similar NSP2 concentration-dependent trend. This suggests very little difference in the overall structure of these segments in the presence or absence of the transcriptome and/or NSP2.

**Figure 6.**
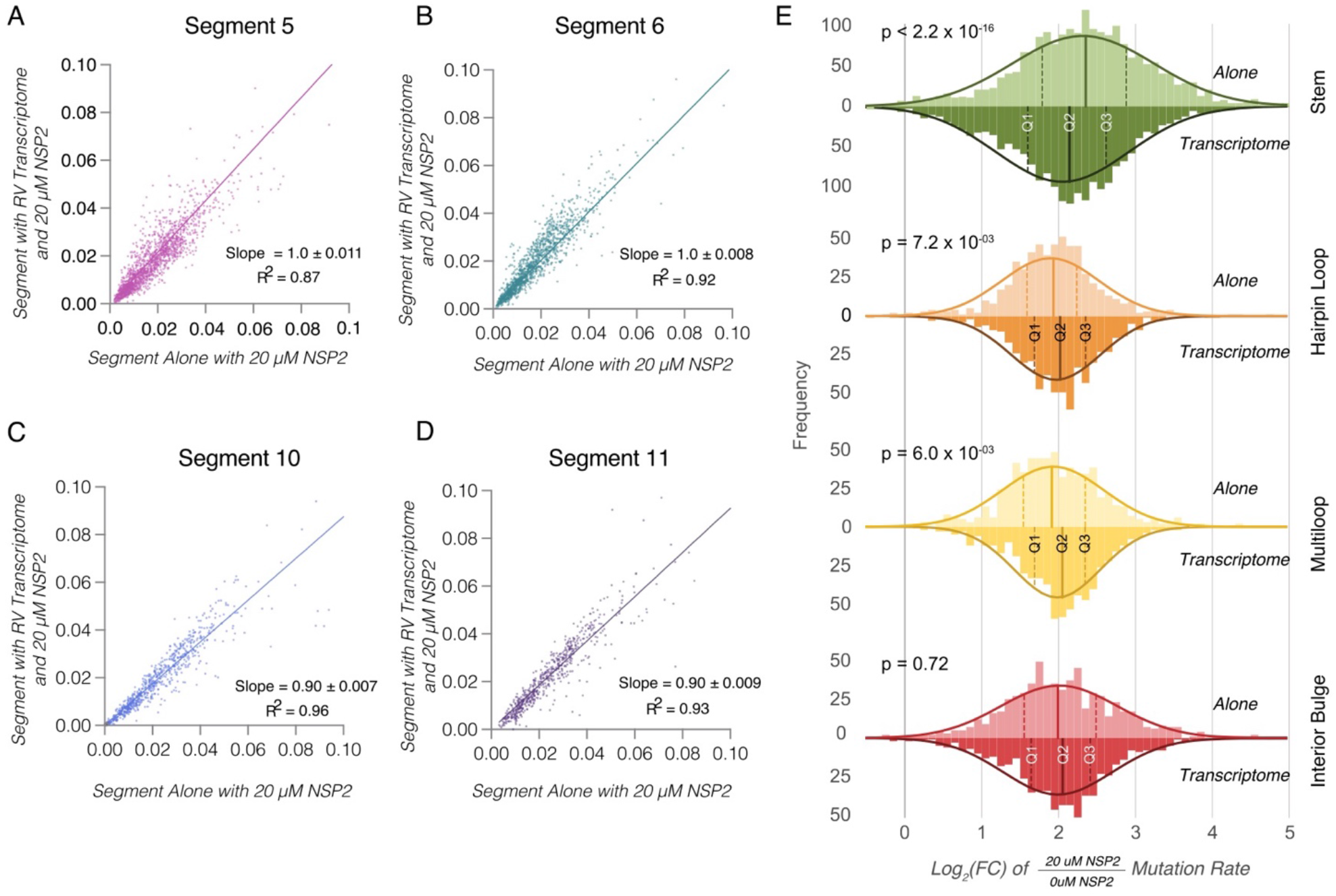
Changes in global backbone flexibility in RV transcriptome mediated by NSP2. Addition of NSP2 to an equimolar mix of 11 distinct RV transcripts (Segment 1 – Segment 11 RNAs constituting RVA transcriptome) results in global increase of mutation rates across all segments. (A-D): Mutation rates of individual RNAs incubated with an RNA mix representing the RV transcriptome in the presence of 20 μM NSP2(y-axis) plotted against mutation rates of the RNA alone in the presence of 20 μM of NSP2 (x-axis) to reveal highly correlated data. (E) Analysis of NSP2-mediated mutation rate changes in the presence or absence of the rotavirus transcriptome. Comparison of distribution of NSP2 mediated mutation rate changes (i.e., the log2 of the quotient of the mutation rates in the presence of 20 μM NSP2 over the mutation rates for RNA without NSP2) for each the structural motif (stem, hairpin loops, multiloops, interior bulge, as described in 5D) for RNAs alone and RNAs in the context of the transcriptome. Q1 and Q3 represent the 25^th^ and 75^th^ interquartile range, respectively, and the median, Q2, is labeled shown as solid lines. Statistical significance was assessed using Kruskal-Wallis test.

**Table 1.** NSP2 increases mutation rates indiscriminately across all RV RNAs. Slopes indicate NSP2 induced increases in mutation rate for each segment calculated by dividing the mutation rate at each point in the titration experiment by the mutation rate of the segment without NSP2 as measured by 5NIA chemical probing. As NPS2 concentration increases, as does the slope. We calculated slope values for the entire RV transcriptome at increasing concentrations of NSP2 and for Segments 5, 6, 10, and 11 alone.

| Segment | 5μM NSP2/0μM NSP2 |  | 10μM NSP2/0μM NSP2 |  | 20μM NSP2/0μM NSP2 |  |
| --- | --- | --- | --- | --- | --- | --- |
|  | Slope | R<sup>2</sup> | Slope | R<sup>2</sup> | Slope | R<sup>2</sup> |
| *Segments probed in the context of RV Transcriptome* |  |  |  |  |  |  |
| Segment 1 | 2.1 ± 0.01 | 0.84 | 3.3 ± 0.03 | 0.67 | 5.4 ± 0.06 | 0.54 |
| Segment 2 | 2.3 ± 0.02 | 0.81 | 3.3 ± 0.04 | 0.70 | 5.4 ± 0.07 | 0.55 |
| Segment 3 | 2.3 ± 0.02 | 0.90 | 3.2 ± 0.03 | 0.79 | 4.5 ± 0.06 | 0.69 |
| Segment 4 | 2.2 ± 0.02 | 0.92 | 3.3 ± 0.03 | 0.81 | 5.5 ± 0.07 | 0.63 |
| Segment 5 | 2.4 ± 0.02 | 0.90 | 3.0 ± 0.03 | 0.88 | 4.2 ± 0.06 | 0.83 |
| Segment 6 | 2.1 ± 0.02 | 0.84 | 2.9 ± 0.05 | 0.73 | 5.9 ± 0.1 | 0.44 |
| Segment 7 | 2.0 ± 0.03 | 0.88 | 2.9 ± 0.05 | 0.79 | 5.7 ± 0.1 | 0.74 |
| Segment 8 | 2.1 ± 0.03 | 0.97 | 2.9 ± 0.04 | 0.96 | 4.9 ± 0.1 | 0.86 |
| Segment 9 | 2.1 ± 0.04 | 0.84 | 3.0 ± 0.04 | 0.79 | 4.2 ± 0.07 | 0.73 |
| Segment 10 | 2.2 ± 0.05 | 0.80 | 2.8 ± 0.08 | 0.65 | 5.9 ± 0.2 | 0.59 |
| Segment 11 | 1.9 ± 0.04 | 0.88 | 2.8 ± 0.08 | 0.73 | 6.6 ± 0.2 | 0.53 |
| *Segments probed alone* |  |  |  |  |  |  |
| Segment 5 | 2.9 ± 0.03 | 0.87 | 2.8 ± 0.03 | 0.88 | 4.8 ± 0.05 | 0.69 |
| Segment 6 | 2.4 ± 0.04 | 0.72 | 3.7 ± 0.06 | 0.56 | 4.8 ± 0.1 | 0.51 |
| Segment 10 | 2.1 ± 0.04 | 0.70 | 3.3 ± 0.08 | 0.42 | 3.9 ± 0.1 | 0.54 |
| Segment 11 | 2.6 ± 0.07 | 0.84 | 4.0 ± 0.1 | 0.67 | 4.6 ± 0.1 | 0.65 |

Remarkably, when we compared the log_2_(FC) distributions for segments 5, 6, 10 and 11 alone and in the presence of the other seven segments (transcriptome), we did observe a structure-specific difference (Figure 6E). What is immediately apparent from the analysis presented in Figure 6E is that the log_2_(FC) is systematically higher for paired nucleotides (as observed in Figure 5E) in both the RNA alone data and transcriptome. However, there are subtle but statistically significant differences in the quartile distributions for paired and unpaired nucleotides. Indeed, the quartile distributions of mutation rates are significantly lower for paired nucleotides (green) in the presence of the transcriptome, while for hairpins and multi-loops they are significantly higher (orange, yellow and red). For interior bulges, the increase is positive, but not significant, likely due to the small number of nucleotides in this conformation relative to loops and stems.

In the context of our previous results, this analysis reveals a subtle but important result that could be explained by transient intermolecular interactions between unpaired nucleotides in the context of the transcriptome. First, neither the addition of NSP2 nor the transcriptome has any detectable effect on the pattern of the SHAPE reactivities (Figure 3A). Thus, it can be concluded that NSP2 is not systematically refolding the RNA in a way that is detectable by chemical probing. Despite fairly subtle changes, our analysis suggests that NSP2 exerts a larger effect on the log_2_(FC) for paired nucleotides.

Remarkably, in the presence of NSP2, paired bases are significantly more reactive towards 5NIA compared to unpaired nucleotides. However, their median reactivity decreases in the presence of the remaining rotaviral transcriptome (Figure 6E, green), consistent with the proposed model that the RNA chaperone NSP2 facilitates inter-molecular RNA-RNA interactions by exposing complementary sequences that are normally sequestered by the local structure (13). Taken together, these data show that quantitative analysis of mutation rates can detect subtle but important structural preferences in promiscuous RNA-binding proteins like NSP2 at transcriptome level, even if they do not appear to have strong structural or sequence preference (41) or when analyzing the normalized SHAPE-MaP data.

## DISCUSSION

In this manuscript we have carried out transcriptome-wide chemical structure probing of eleven rotavirus A transcripts that serve as RV genome precursors. Our data validates previous computational and co-variance models of some of the RV RNAs, providing experimental support for these structures. Importantly, we have identified multiple well-defined structured regions of low SHAPE reactivity across the entire transcriptome (Figure 1B). Thus, our data suggest that RNA structure is pervasive in rotaviral transcripts, and it is not limited to untranslated terminal regions. Pervasive RNA secondary structures is a feature of all RNA viruses studied to date using chemical probing including flaviviruses (42), H1N1 (36), HIV (9), alphaviruses (10, 34), and coronaviruses (43, 44). It must be noted that the structures we have identified here only exist prior to genome packaging (1, 45, 46). Nonetheless, we observe globally similarly sized regions of low SHAPE as in other RNA viruses (ranging from 50-200 nucleotides), suggesting these pervasive secondary structures tend to emerge in most large viral RNAs. We were interested to use chemical structure probing to evaluate how NSP2, a key RNA chaperone involved in RNA assortment and packaging may interact with these pervasive structures.

Despite multiple biophysical and biochemical studies that have demonstrated helix-destabilizing activity of NSP2 with shorter RNA substrates (12, 13, 15), surprisingly, incubation of any of the rotaviral segments with even large molar excesses of NSP2 had no significant effect on SHAPE reactivity. At first, these results appeared to be at odds with our previous biophysical studies of the RNA chaperoning effect of NSP2 on small RNA stem-loops that revealed melting of the structure (13, 15). However, our direct analysis of adduct-induced mutations via mutational profiling allowed us to observe the previously unknown global effect of NSP2 on the rotaviral transcripts. Indeed, the quartile normalization used in the SHAPE-MaP data processing effectively masks any large global changes in mutation rates. Thus, although next generation sequencing enables unprecedented throughput, detailed understanding of the experimental system (i.e., specific vs non-specific RNA binding, RNA:protein binding stoichiometry, complementary biophysical data supporting RNA conformational changes) should always be considered for careful SHAPE-MaP data interpretation. Furthermore, visualization of the raw mutational data during analysis should always be carried out to reveal any changes in the RNA backbone dynamics that might be normalized out in subsequent data processing steps.

Having established that we observe a very significant, global, and concentration-dependent increase in mutation rate upon addition of NSP2, we decided to evaluate whether we could observe any structural specificity. It has been suggested previously that with small RNAs, NSP2 preferentially binds unpaired nucleotides with sub-nanomolar affinity (13). Our classification analysis of bases as paired or unpaired nucleotides allowed us to observe that the log_2_(Fold Change) in mutation rate upon addition of NSP2 was more positive for paired nucleotides across entire RV transcriptome (Figure 6). Previous studies suggest that NSP2 preferentially binds unpaired RNA regions (41), therefore the higher log_2_(Fold Change) for paired regions does not reflect any structural preference in binding of NSP2. Rather, our mutational profiling analysis indicates that NSP2 has a larger relative effect on paired nucleotide’s flexibility, consistent with its role as an RNA chaperone that promotes RNA-RNA interactions by making sequestered complementary sequences more accessible for making new inter-molecular contacts. It should be noted that further in-depth analyses followed by an extensive functional validation would be required to identify the nucleotides involved in the assembly of eleven distinct RV transcripts. Recent studies of the assortment and packaging of 8 RNA segments in influenza A viruses suggest that in IAVs such interactions are likely to be highly redundant, and potentially transient in nature (36, 47), thus making their identification and validation a particularly challenging task.

Nevertheless, understanding the mechanistic role of promiscuous RNA-binding proteins with RNA chaperone activities will help our endeavors to elucidate the RNA assembly pathways leading to a stoichiometric segment packaging. It should be noted that in RV-infected cells, based on the RNA-Seq estimates, the cytoplasmic viral transcript concentration is around 10-100 nM (48), while the intracellular NSP2 concentration reaches 1-10 μM or higher (25). Thus, our *in vitro* assays are broadly in line with the physiologically relevant RNA:NSP2 molar ratios that also promote RNA:RNA interactions *in vitro* (13).

It is important to note that the increase in mutation rate indicates a global change in RNA backbone flexibility, but does not change the overall SHAPE pattern, which informs RNA structure. Thus, one should not interpret this increase as an overall melting of secondary structure, or rearrangement of base-pairing. Indeed, we performed an analogous NSP2 titration using dimethyl sulfate (DMS), which methylates unpaired adenines and cytosines, and does not affect the backbone. We did not observe a significant increase in mutation rate, consistent with the base-pairing remaining largely unperturbed by NSP2 (Supplementary Figure S4). Thus, the DMS probing data and corresponding mutation rate analysis support our hypothesis that NSP2 binding increases RNA backbone flexibility, while having very little effect on Watson-Crick base-pairing under our experimental conditions. It should be noted that SHAPE reagents measure backbone flexibility by preferentially acylating 2' hydroxyls adopting a specific flexible geometry (22, 33). It is the relationship between backbone flexibility and base-pairing that allows us to use SHAPE reactivity to inform secondary structure prediction, but fundamentally SHAPE reagents measure flexibility. Thus, the observed global mutation rate increase for SHAPE (but not DMS) indicates the effect of NSP2 is limited to the RNA backbone.

These data further support a model in which NSP2 is able to capture folded RNA stem-loop structures within a single RNA-binding groove, resulting in general relaxation of RNA structure without fraying of the RNA stem termini (15). This allows the RNA structure to be sufficiently flexible so that it can adopt a conformation compatible with stabilization of sequence-specific, potentially transient RNA-RNA contacts. By binding to multiple RNAs concurrently via surface-exposed positively charged grooves, NSP2 octamers act as matchmakers of complementary sequences, promoting intermolecular RNA–RNA interactions (15). While NSP2 promiscuously binds any ssRNA with sub-nanomolar affinity, the proposed mechanism would require NSP2-RNA interactions to be transient to allow sampling of multiple interacting RNA partners. We propose that on a transcriptome level, NSP2 achieves its RNA matchmaking activity by globally increasing RNA backbone flexibility, while the formation of multiple regions of inter-molecular base-pairing is governed by specific, yet to be revealed, complementary sequences present in the RV transcriptome.

## REFERENCES

1. Gallegos, C.O. and Patton, J.T. (1989) Characterization of rotavirus replication intermediates: A model for the assembly of single-shelled particles. Virology, 172, 616–627.

2. Jayaram, H., Estes, M.K. and Prasad, B.V.V. (2004) Emerging themes in rotavirus cell entry, genome organization, transcription and replication. Virus Res., 101, 67–81.

3. McDonald, S.M. and Patton, J.T. (2011) Assortment and packaging of the segmented rotavirus genome. Trends Microbiol., 19, 136–144.

4. Silvestri, L.S., Taraporewala, Z.F. and Patton, J.T. (2004) Rotavirus Replication: Plus-Sense Templates for Double-Stranded RNA Synthesis Are Made in Viroplasms. J. Virol., 78, 7763–7774.

5. Li, W., Manktelow, E., von Kirchbach, J.C., Gog, J.R., Desselberger, U. and Lever, A.M. (2010) Genomic analysis of codon, sequence and structural conservation with selective biochemical-structure mapping reveals highly conserved and dynamic structures in rotavirus RNAs with potential cis-acting functions. Nucleic Acids Res., 38, 7718–7735.

6. Poncet, D., Aponte, C. and Cohen, J. (1993) Rotavirus protein NSP3 (NS34) is bound to the 3’ end consensus sequence of viral mRNAs in infected cells. J. Virol., 67, 3159–3165.

7. Vende, P., Piron, M., Castagné, N. and Poncet, D. (2000) Efficient Translation of Rotavirus mRNA Requires Simultaneous Interaction of NSP3 with the Eukaryotic Translation Initiation Factor eIF4G and the mRNA 3′ End. J. Virol., 74, 7064–7071.

8. Desselberger, U. (2017) Reverse genetics of rotavirus. Proc. Natl. Acad. Sci., 114, 2106–2108.

9. Watts, J.M., Dang, K.K., Gorelick, R.J., Leonard, C.W., Bess, J.W., Swanstrom, R., Burch, C.L. and Weeks, K.M. (2009) Architecture and Secondary Structure of an Entire HIV-1 RNA Genome. Nature, 460, 711–716.

10. Madden, E.A., Plante, K.S., Morrison, C.R., Kutchko, K.M., Sanders, W., Long, K.M., Taft-Benz, S., Cruz Cisneros, M.C., White, A.M., Sarkar, S., et al. Using SHAPE-MaP To Model RNA Secondary Structure and Identify 3′UTR Variation in Chikungunya Virus. J. Virol., 94, e00701–20.

11. Alexander Borodavka, Ulrich Desselberger, Ulrich Desselberger, and John T. Patton (2018) Genome packaging in multi-segmented dsRNA viruses: distinct mechanisms with similar outcomes. Curr. Opin. Virol., 10.1016/j.coviro.2018.08.001.

12. Taraporewala, Z.F. and Patton, J.T. (2001) Identification and Characterization of the Helix-Destabilizing Activity of Rotavirus Nonstructural Protein NSP2. J. Virol., 75, 4519–4527.

13. Borodavka, A., Dykeman, E.C., Schrimpf, W. and Lamb, D.C. (2017) Protein-mediated RNA folding governs sequence-specific interactions between rotavirus genome segments. eLife, 6, e27453.

14. Woodson, S.A. (2010) Taming free energy landscapes with RNA chaperones. RNA Biol., 7, 677–686.

15. Bravo, J.P.K., Bartnik, K., Venditti, L., Acker, J., Gail, E.H., Colyer, A., Davidovich, C., Lamb, D.C., Tuma, R., Calabrese, A.N., et al. (2021) Structural basis of rotavirus RNA chaperone displacement and RNA annealing. Proc. Natl. Acad. Sci., 118.

16. Warf, M.B. and Berglund, J.A. (2010) Role of RNA structure in regulating pre-mRNA splicing. Trends Biochem. Sci., 35, 169–178.

17. Mortimer, S.A., Kidwell, M.A. and Doudna, J.A. (2014) Insights into RNA structure and function from genome-wide studies. Nat. Rev. Genet., 15, 469–479.

18. Deigan, K.E., Li, T.W., Mathews, D.H. and Weeks, K.M. (2009) Accurate SHAPE-directed RNA structure determination. Proc. Natl. Acad. Sci., 106, 97–102.

19. Merino, E.J., Wilkinson, K.A., Coughlan, J.L. and Weeks, K.M. (2005) RNA Structure Analysis at Single Nucleotide Resolution by Selective 2’-Hydroxyl Acylation and Primer Extension (SHAPE). J. Am. Chem. Soc., 127, 4223–4231.

20. Nicholson, B.L. and White, K.A. (2014) Functional long-range RNA–RNA interactions in positive-strand RNA viruses. Nat. Rev. Microbiol., 12, 493–504.

21. Tuplin, A., Struthers, M., Simmonds, P. and Evans, D.J. (2012) A twist in the tail: SHAPE mapping of long-range interactions and structural rearrangements of RNA elements involved in HCV replication. Nucleic Acids Res., 40, 6908–6921.

22. Smola, M.J., Rice, G.M., Busan, S., Siegfried, N.A. and Weeks, K.M. (2015) Selective 2′-hydroxyl acylation analyzed by primer extension and mutational profiling (SHAPE-MaP) for direct, versatile and accurate RNA structure analysis. Nat. Protoc., 10, 1643–1669.

23. Richards, J.E., Desselberger, U. and Lever, A.M. (2013) Experimental Pathways towards Developing a Rotavirus Reverse Genetics System: Synthetic Full Length Rotavirus ssRNAs Are Neither Infectious nor Translated in Permissive Cells. PLoS ONE, 8, e74328.

24. Borodavka, A., Singaram, S.W., Stockley, P.G., Gelbart, W.M., Ben-Shaul, A. and Tuma, R. (2016) Sizes of Long RNA Molecules Are Determined by the Branching Patterns of Their Secondary Structures. Biophys. J., 111, 2077–2085.

25. Geiger, F., Acker, J., Papa, G., Wang, X., Arter, W.E., Saar, K.L., Erkamp, N.A., Qi, R., Bravo, J.P., Strauss, S., et al. (2021) Liquid–liquid phase separation underpins the formation of replication factories in rotaviruses. EMBO J., 40, e107711.

26. Tijerina, P., Mohr, S. and Russell, R. (2007) DMS footprinting of structured RNAs and RNA-protein complexes. Nat. Protoc., 2, 2608–2623.

27. Busan, S. and Weeks, K.M. (2018) Accurate detection of chemical modifications in RNA by mutational profiling (MaP) with ShapeMapper 2. RNA N. Y. N, 24, 143–148.

28. Low, J.T. and Weeks, K.M. (2010) SHAPE-Directed RNA Secondary Structure Prediction. Methods San Diego Calif, 52, 150–158.

29. Reuter, J.S. and Mathews, D.H. (2010) RNAstructure: software for RNA secondary structure prediction and analysis. BMC Bioinformatics, 11, 129.

30. Thiel, B.C., Beckmann, I.K., Kerpedjiev, P. and Hofacker, I.L. (2019) 3D based on 2D: Calculating helix angles and stacking patterns using forgi 2.0, an RNA Python library centered on secondary structure elements. F1000Research, 8, ISCB Comm J–287.

31. Wickham, H. (2016) ggplot2: Elegant Graphics for Data Analysis Springer-Verlag New York.

32. Busan, S., Weidmann, C.A., Sengupta, A. and Weeks, K.M. (2019) Guidelines for SHAPE Reagent Choice and Detection Strategy for RNA Structure Probing Studies. Biochemistry, 58, 2655–2664.

33. Siegfried, N.A., Busan, S., Rice, G.M., Nelson, J.A.E. and Weeks, K.M. (2014) RNA motif discovery by SHAPE and mutational profiling (SHAPE-MaP). Nat. Methods, 11, 959–965.

34. Kutchko, K.M., Madden, E.A., Morrison, C., Plante, K.S., Sanders, W., Vincent, H.A., Cruz Cisneros, M.C., Long, K.M., Moorman, N.J., Heise, M.T., et al. (2018) Structural divergence creates new functional features in alphavirus genomes. Nucleic Acids Res., 46, 3657–3670.

35. Smola, M.J. and Weeks, K.M. (2018) In-cell RNA structure probing with SHAPE-MaP. Nat. Protoc., 13, 1181–1195.

36. Dadonaite, B., Gilbertson, B., Knight, M.L., Trifkovic, S., Rockman, S., Laederach, A., Brown, L.E., Fodor, E. and Bauer, D.L.V. (2019) The structure of the influenza A virus genome. Nat. Microbiol., 4, 1781–1789.

37. Woods, C.T., Lackey, L., Williams, B., Dokholyan, N.V., Gotz, D. and Laederach, A. (2017) Comparative Visualization of the RNA Suboptimal Conformational Ensemble In Vivo. Biophys. J., 113, 290–301.

38. Lackey, L., Coria, A., Woods, C., McArthur, E. and Laederach, A. (2018) Allele-specific SHAPE-MaP assessment of the effects of somatic variation and protein binding on mRNA structure. RNA N. Y. N, 24, 513–528.

39. Yang, A.-D., Barro, M., Gorziglia, M.I. and Patton, J.T. (2004) Translation enhancer in the 3?-untranslated region of rotavirus gene 6 mRNA promotes expression of the major capsid protein VP6. Arch. Virol., 149, 303–321.

40. Gog, J.R., Afonso, E.D.S., Dalton, R.M., Leclercq, I., Tiley, L., Elton, D., von Kirchbach, J.C., Naffakh, N., Escriou, N. and Digard, P. (2007) Codon conservation in the influenza A virus genome defines RNA packaging signals. Nucleic Acids Res., 35, 1897–1907.

41. Bravo, J.P.K., Borodavka, A., Barth, A., Calabrese, A.N., Mojzes, P., Cockburn, J.J.B., Lamb, D.C. and Tuma, R. (2018) Stability of local secondary structure determines selectivity of viral RNA chaperones. Nucleic Acids Res., 46, 7924–7937.

42. Dethoff, E.A., Boerneke, M.A., Gokhale, N.S., Muhire, B.M., Martin, D.P., Sacco, M.T., McFadden, M.J., Weinstein, J.B., Messer, W.B., Horner, S.M., et al. (2018) Pervasive tertiary structure in the dengue virus RNA genome. Proc. Natl. Acad. Sci., 115, 11513–11518.

43. Schlick, T., Zhu, Q., Dey, A., Jain, S., Yan, S. and Laederach, A. (2021) To Knot or Not to Knot: Multiple Conformations of the SARS-CoV-2 Frameshifting RNA Element. J. Am. Chem. Soc., 143, 11404–11422.

44. Huston, N.C., Wan, H., Strine, M.S., de Cesaris Araujo Tavares, R., Wilen, C.B. and Pyle, A.M. (2021) Comprehensive in vivo secondary structure of the SARS-CoV-2 genome reveals novel regulatory motifs and mechanisms. Mol. Cell, 81, 584-598.e5.

45. Acs, G., Klett, H., Schonberg, M., Christman, J., Levin, D.H. and Silverstein, S.C. (1971) Mechanism of Reovirus Double-Stranded Ribonucleic Acid Synthesis In Vivo and In Vitro. J. Virol., 8, 684– 689.

46. Desselberger, U., Richards, J., Tchertanov, L., Lepault, J., Lever, A., Burrone, O. and Cohen, J. (2013) Further characterisation of rotavirus cores: Ss(+)RNAs can be packaged in vitro but packaging lacks sequence specificity. Virus Res., 178, 252–263.

47. Haralampiev, I., Prisner, S., Nitzan, M., Schade, M., Jolmes, F., Schreiber, M., Loidolt-Krüger, M., Jongen, K., Chamiolo, J., Nilson, N., et al. (2020) Selective flexible packaging pathways of the segmented genome of influenza A virus. Nat. Commun., 11, 4355.

48. Strauss, S., Borodavka, A., Papa, G., Desiró, D., Schueder, F. and Jungmann, R. (2021) Principles of RNA recruitment to viral ribonucleoprotein condensates in a segmented dsRNA virus. bioRxiv, 10.1101/2021.03.22.435476.

