## Supporting Data for "Rotavirus RNA chaperone mediates global transcriptome-wide increase in RNA backbone flexibility"

A

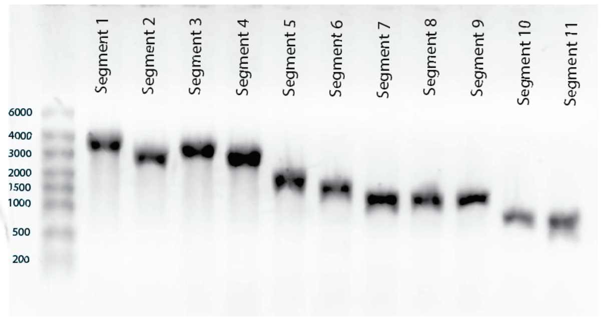


B

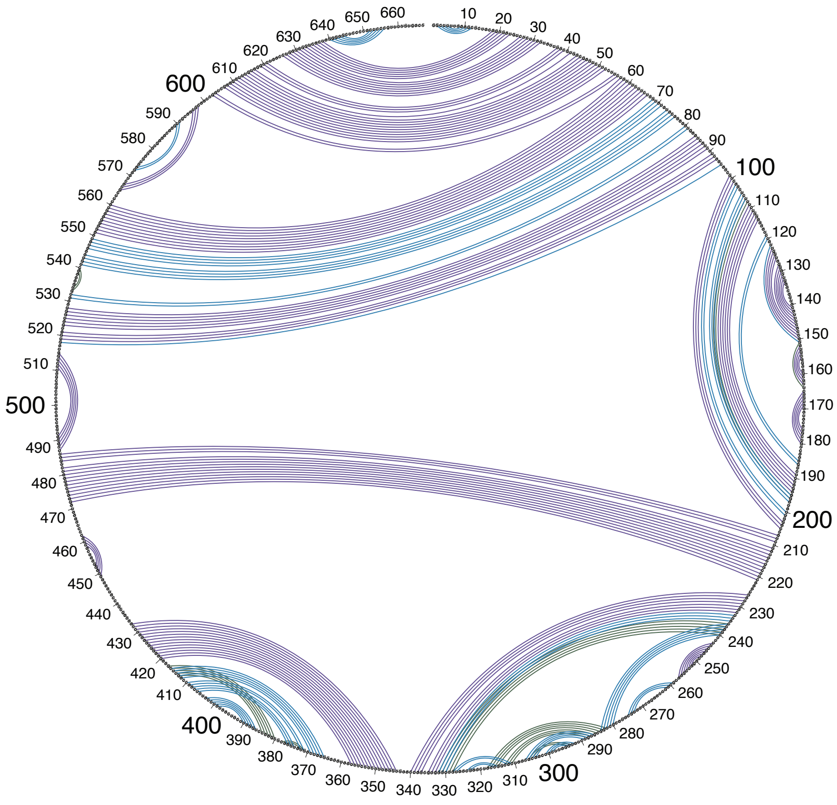


**Supplementary Figure S1.** Rotavirus RNA templates used in this study and comparison of previously predicted segment 11 secondary structure with our experimentally validated segment 11 secondary structure. (A) In vitro synthesized RV segments 1- 11. The RNAs were synthesized from digested plasmid products, coding for RF segments (Materials and Methods). 1 µg of each RNA were resolved on a denaturing MOPS gel (1% agarose, 1X MOPS, 7.2% formaldehyde). These RNAs were then aliquoted, stored in -80, and used for the studies described in this paper. (B) Circle comparison of our SHAPE informed secondary structure diagram of Segment 11 and a SELEX informed segment 11 secondary structure prediction we previously proposed (Borodavka et al. 2017; Reuter and Mathews 2010). Individual lines in the circle represents a predicted base pair, base pairs in purple represent base pairs that both secondary structure models share, blue lines are base pairs only present in the SHAPE-MaP derived secondary structure we present here, and green lines represent base pairs in the SELEX informed secondary structure model. We found that both models were broadly similar with a PPV of 66.8% and sensitivity of 84.52%.

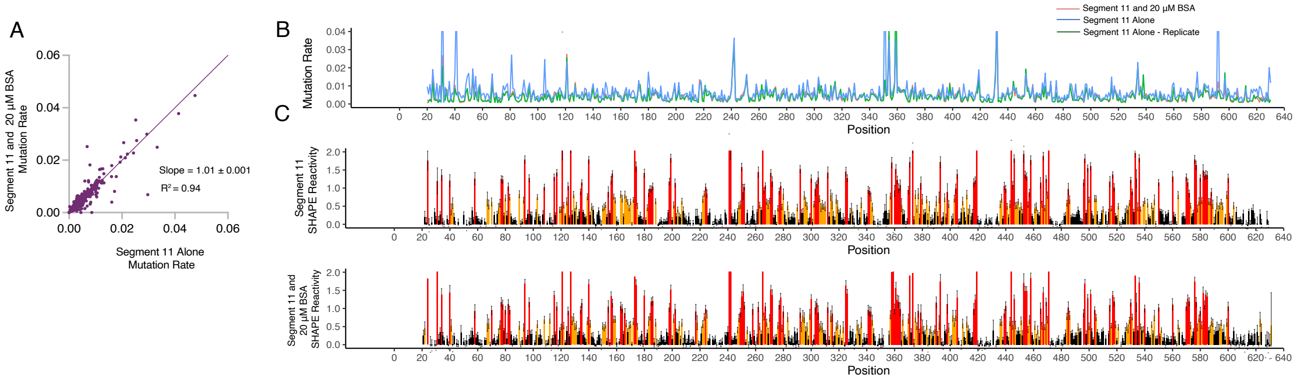


**Supplementary Figure S2.** BSA has no impact on Segment 11 secondary structure or mutation rates.
(A) Scatter plot comparing mutation rate of Segment 11 alone and Segment 11 and 20 µM NSP2, the data sets are identical as measured by the slope between the two data sets (1.0 ± 0.001) and $R^2$ = 0.0.94. (B) Linear representation of the mutation rate data in the scatter plot across the Segment 11 transcript. Red represents the mutation rate of Segment 11 with 5NIA and 20 µM BSA, the blue line represents segment 11 alone, and the green line represents a replicate of Segment 11 alone. (C) Normalized SHAPE reactivities of the mutation rates shown in panel B. Top: SHAPE reactivity profile of Segment 11 RNA alone. Bottom: SHAPE reactivity profile of Segment 11 incubated with 20 µM NSP2

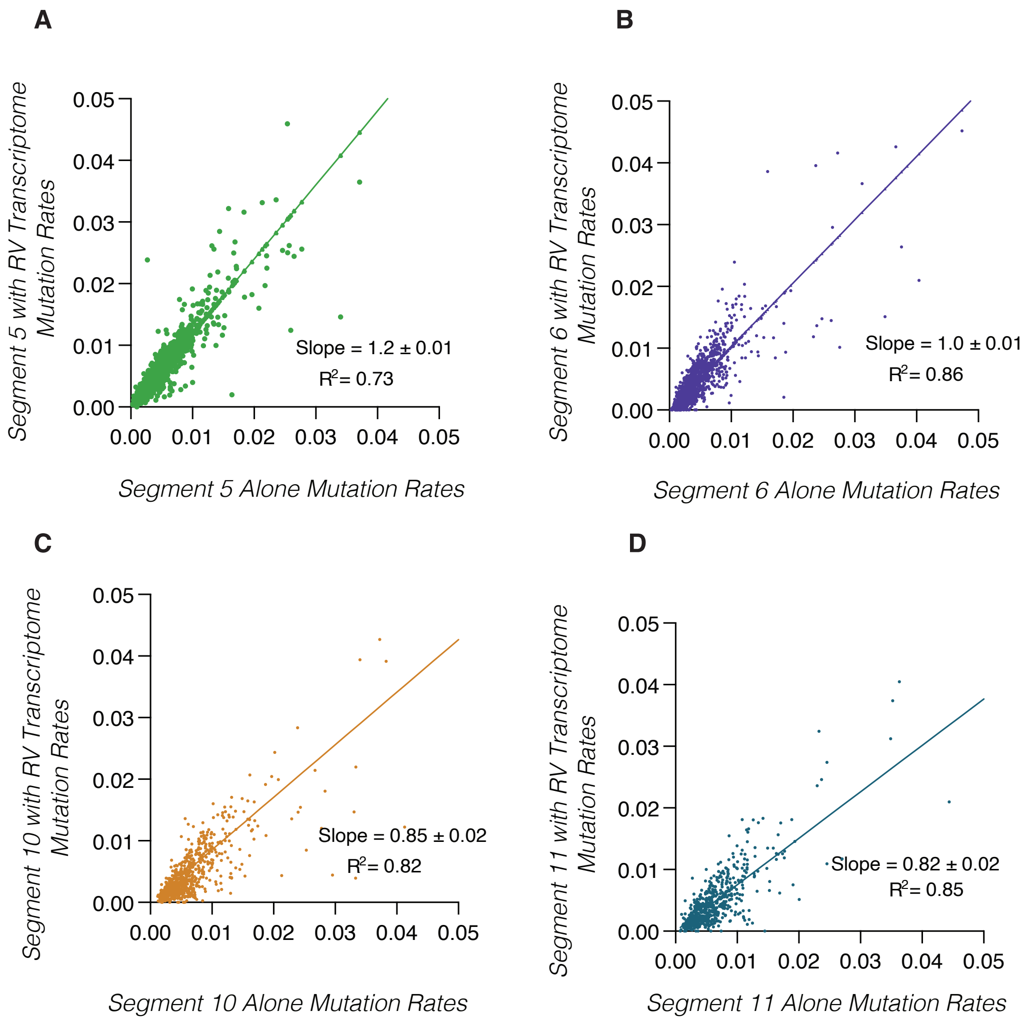


**Supplementary Figure S3.** Presence of RV RNAs doesn't significantly change segment mutation rate.

(A-D) Scatter plot comparing the mutation rate data of Segments 5, 6, 10, 11 alone (x axis) with the mutation rates of the same segments in the context of the entire transcriptome (y axis). The data sets are highly similar with slopes around 1 and high correlations $R^2$ close to 1.

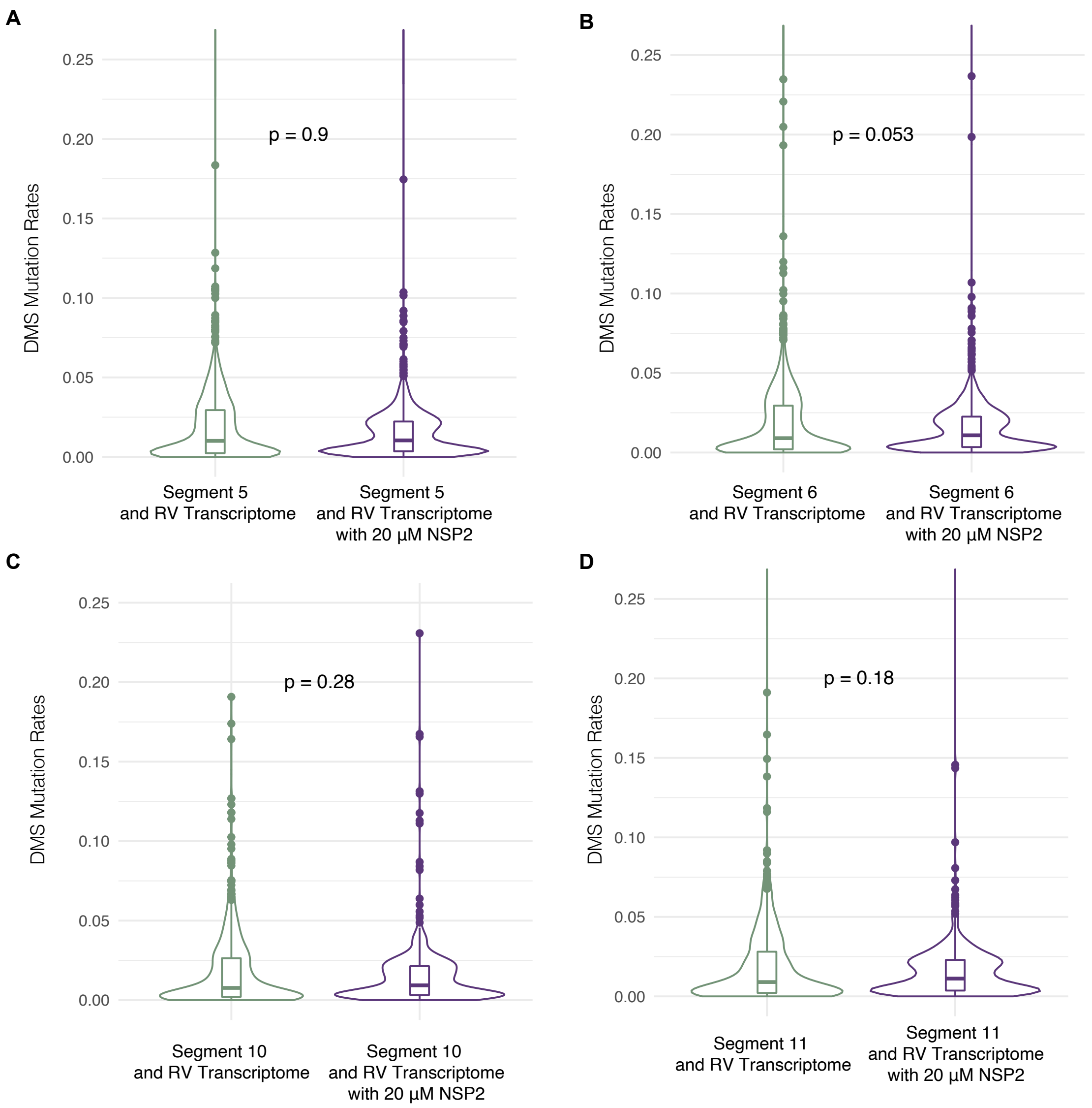


**Supplementary Figure S4.** NSP2 does not change Segment 5, 6, 10, and 11 DMS mutation rates. (A-D) Violin plots comparing distribution of DMS induced mutation rates of denoted RV segment when incubated in the presence of all RV Segments or in the presence of both all RV Segments and 20 µM NSP2. DMS methylates the Watson-Crick base-pairing edge of unpaired adenine and cytosine bases. Boxes represent the 25th/75th interquartile range, and medians are shown as central bands. Significance values were calculated using Kruskal-Wallis test ($p < 0.05$)

| | Rep 1 vs Rep 2: 0 µM NSP2 | | Rep 1 vs Rep 2: 5 µM NSP2 | | Rep 1 vs Rep 2: 10 µM NSP2 | | Rep 1 vs Rep 2: 20 µM NSP2 | |
|---|---|---|---|---|---|---|---|---|
| Segment | Slope | $R^2$ | Slope | $R^2$ | Slope | $R^2$ | Slope | $R^2$ |
| *Segments probed in the context of RV Transcriptome* | | | | | | | | |
| Segment 1 | 0.97 ± 0.003 | 0,97 | 1.1 ± 0.003 | 0,97 | 0.95 ± 0.003 | 0,97 | 0.80 ± 0.003 | 0,97 |
| Segment 2 | 0.98 ± 0.003 | 0,97 | 1.0 ± 0.003 | 0,97 | 0.95 ± 0.004 | 0,96 | 0.80 ± 0.003 | 0,97 |
| Segment 3 | 0.97 ± 0.004 | 0,96 | 1.0 ± 0.004 | 0,97 | 0.95 ± 0.003 | 0,98 | 0.93 ± 0.003 | 0,98 |
| Segment 4 | 1.0 ± 0.01 | 0,79 | 1.0 ± 0.01 | 0,93 | 0.98 ± 0.01 | 0,90 | 0.92 ± 0.01 | 0,94 |
| Segment 5 | 1.0 ± 0.01 | 0,92 | 1.0 ± 0.004 | 0,97 | 0.96 ± 0.003 | 0,98 | 0.96 ± 0.004 | 0,98 |
| Segment 6 | 0.99+0.01 | 0,95 | 1.1 ± 0.01 | 0,96 | 1.0 ± 0.01 | 0,85 | 0.98 ± 0.01 | 0,96 |
| Segment 7 | 0.94+0.01 | 0,97 | 1.1 ± 0.01 | 0,97 | 0.93 ± 0.01 | 0,98 | 0.96 ± 0.003 | 0,99 |
| Segment 8 | 0.95+0.01 | 0,94 | 1.0 ± 0.01 | 0,96 | 0.95 ± 0.01 | 0,96 | 0.95 ± 0.01 | 0,98 |
| Segment 9 | 0.95+0.01 | 0,97 | 1.1 ± 0.01 | 0,98 | 0.95 ± 0.004 | 0,98 | 0.95 ± 0.004 | 0,98 |
| Segment 10 | 0.98 ± 0.01 | 0,93 | 0.90 ± 0.01 | 0,96 | 0.95 ± 0.01 | 0,93 | 0.98 ± 0.01 | 0,93 |
| Segment 11 | 1.1 ± 0.06 | 0,98 | 0.96 ± 0.07 | 0,97 | 0.89 ± 0.01 | 0,94 | 1.1 ± 0.01 | 0,96 |
| *Segments probed alone* | | | | | | | | |
| Segment 5 | 1.3 ± 0.01 | 0,89 | 0.93 ± 0.01 | 0,84 | 1.0 ± 0.02 | 0,80 | 1.0 ± 0.01 | 0,94 |
| Segment 6 | 0.98 ± 0.02 | 0,68 | 1.0 ± 0.01 | 0,90 | 1.2 ± 0.01 | 0,93 | 1.1 ± 0.01 | 0,91 |
| Segment 10 | 1.0 ± 0.01 | 0,98 | 0.90 ± 0.004 | 0,99 | 0.94 ± 0.004 | 0,99 | 0.94 ± 0.003 | 0,99 |
| Segment 11 | 1.1 ± 0.03 | 0,74 | 1.1 ± 0.02 | 0,89 | 0.98 ± 0.01 | 0,98 | 0.99 ± 0.01 | 0,98 |

**Supplementary Table S1.** Similarity and correlation between mutation rates of data sets of 11 segments all together and of segments 5, 6, 10, and 11 alone.

| | Rep 1 vs Rep 2: 0 µM NSP2 | | Rep 1 vs Rep 2: 5 µM NSP2 | | Rep 1 vs Rep 2: 10 µM NSP2 | | Rep 1 vs Rep 2: 20 µM NSP2 | |
|---|---|---|---|---|---|---|---|---|
| Segment | Slope | $R^2$ | Slope | $R^2$ | Slope | $R^2$ | Slope | $R^2$ |
| *Segments probed in the context of RV Transcriptome* | | | | | | | | |
| Segment 1 | 0.96 ± 0.01 | 0,91 | 0.92 ± 0.01 | 0,91 | 0.97 ± 0.003 | 0,97 | 1.0 ± 0.003 | 0,96 |
| Segment 2 | 0.94 ± 0.01 | 0,91 | 0.93 ± 0.01 | 0,93 | 0.96 ± 0.004 | 0,95 | 1.0 ± 0.004 | 0,96 |
| Segment 3 | 0.90 ± 0.01 | 0,87 | 0.87 ± 0.004 | 0,93 | 1.0 ± 0.004 | 0,97 | 1.0 ± 0.004 | 0,97 |
| Segment 4 | 0.96 ± 0.01 | 0,91 | 0.92 ± 0.02 | 0,91 | 0.98 ± 0.004 | 0,97 | 1.0 ± 0.004 | 0,97 |
| Segment 5 | 0.92 ± 0.01 | 0,92 | 0.94 ± 0.01 | 0,90 | 1.0 ± 0.004 | 0,97 | 0.96 ± 0.004 | 0,97 |
| Segment 6 | 0.90 ± 0.01 | 0,78 | 0.85 ± 0.01 | 0,82 | 0.97 ± 0.01 | 0,95 | 0.92 ± 0.004 | 0,97 |
| Segment 7 | 0.94 ± 0.01 | 0,88 | 0.88 ± 0.01 | 0,88 | 1.0 ± 0.01 | 0,97 | 1.0 ± 0.004 | 0,98 |
| Segment 8 | 0.90 ± 0.01 | 0,80 | 0.89 ± 0.01 | 0,85 | 0.98 ± 0.01 | 0,95 | 0.96 ± 0.01 | 0,97 |
| Segment 9 | 0.90 ± 0.01 | 0,88 | 0.89 ± 0.01 | 0,90 | 0.98 ± 0.01 | 0,96 | 0.96 ± 0.01 | 0,97 |
| Segment 10 | 0.86 ± 0.02 | 0,73 | 0.80 ± 0.02 | 0,77 | 0.97 ± 0.01 | 0,92 | 0.92 ± 0.01 | 0,97 |
| Segment 11 | 0.93 ± 0.02 | 0,80 | 0.86 ± 0.02 | 0,83 | 0.97 ± 0.01 | 0,92 | 0.90 ± 0.01 | 0,97 |
| *Segments probed alone* | | | | | | | | |
| Segment 5 | 0.89 ± 0.01 | 0,79 | 1.0 ± 0.02 | 0,68 | 0.93 ± 0.01 | 0,78 | 0.95 ± 0.01 | 0,93 |
| Segment 6 | 0.84 ± 0.02 | 0,64 | 1.0 ± 0.01 | 0,94 | 1.0 ± 0.01 | 0,95 | 0.98 ± 0.01 | 0,92 |
| Segment 10 | 1.0 ± 0.01 | 0,97 | 1.0 ± 0.004 | 0,99 | 1.0 ± 0.004 | 0,99 | 1.0 ± 0.003 | 0,99 |
| Segment 11 | 0.94 ± 0.02 | 0,93 | 0.97 ± 0.01 | 0,96 | 0.99 ± 0.003 | 0,99 | 0.96 ± 0.01 | 0,99 |

**Supplementary Table S2** Similarity and correlation between SHAPE-MaP reactivities of data sets of 11 segments all together and of segments 5, 6, 10, and 11 alone.

**Supplementary File S3** All SHAPE data used in this study in SNRNASM format.
